# How climate change and population growth will shape attendance and human-wildlife interactions at British Columbia parks

**DOI:** 10.1101/2023.07.11.548618

**Authors:** Dayna K Weststrate, Aimee Chhen, Stefano Mezzini, Kirk Safford, Michael J Noonan

## Abstract

Protected areas are important for ecological conservation while simultaneously supporting culturally, and economically valuable tourism. However, excessive guest volumes strain operations and risk human-wildlife conflict, threatening the sustainability of nature-based tourism. Thus, park managers need to know what factors underpin attendance and how these might interact to shape future attendance. Using a decade of attendance records from 249 provincial parks, in British Columbia (BC), Canada, as well as 12 years of human-wildlife interactions (HWI) records at five national parks in BC, we modelled the impacts of weather conditions and population growth interact on park attendance and HWIs. We paired these models with climate change and population growth scenarios to generate projections of how attendance and HWIs will change throughout the century. Climate change is projected to result in more precipitation and higher temperatures, and, over this same time span, BC’s population is expected to grow substantially. Based on the observed relationship between attendance and weather, parks should anticipate a marked rise in visitors and HWIs especially during their respective peak seasons. These projections provide park managers with the information required for proactive management, ultimately contributing to the sustainability of recreation and tourism in protected areas.

## Introduction

Conservation, tourism, and outdoor recreation have developed a long history of working in tandem through protected areas toward mutually beneficial outcomes (Stronza et al., 2019). Protected areas are essential for conservation: they play an important role in protecting local flora and fauna, and they promote high species richness and biodiversity (Bass et al., 2010; C. Chen et al., 2022; Gray et al., 2016; Margules & Pressey, 2000; Thomas & Gillingham, 2015). In turn, many protected areas serve as culturally and economically important tourism and recreation sites, offering significant economic value by way of employment (Eagles, 2002), GDP contribution (Buckley, 2018), and the bolstering of local economies (Sangpikul, 2017; Taylor et al., 2003). Indeed, the mental health benefits of visiting protected areas alone have been valued at US$6 trillion (Buckley et al., 2019). Yet, even though biodiverse protected areas can reinforce tourism benefits and vice versa (Stronza et al., 2019), nearly 40% of protected areas worldwide are operating under severe human pressure (Jones et al., 2018).

Guest volume is a pressing issue for park managers, as it strains operations and reduces the quality of the visitor experience (Manning, 2001; Prakash et al., 2019). The presence and behaviour of humans in protected areas can negatively impact wildlife through increased zoonotic disease transmission (Charron, 2002; Monahan et al., 2009), a loss of biodiversity (Nyhus, 2016), altered spatiotemporal patterns in habitat use, including habitat loss through avoidance, behavioural change (e.g., conditioning, tolerance, predator shield; Lopez Gutierrez et al., 2020; Procko et al., 2023), and changes in energetics (Corradini et al., 2021; Gaynor et al., 2018; Larson et al., 2016; Reed & Merenlender, 2008; Rogala et al., 2011; Sarmento & Berger, 2017; Whittington et al., 2022). Human sourced food and salts also act as attractants for many species, generating human-wildlife-interactions (HWIs) that can result in the destruction of individual animals (Hebblewhite et al., 2003; Vayro et al., 2023). Beyond the impacts on wildlife, HWIs also pose a risk of physical endangerment to park visitors, property damage, and a loss of recreational opportunities (Nyhus, 2016). As protected areas receive more guests, these issues become amplified and HWIs become difficult to avoid, resulting in more frequent and severe conflict (Cui et al., 2021; Geng, 2021). The managers of protected areas thus often find themselves on the frontlines of socioecologically challenging human-wildlife-conflict issues. If HWIs cannot be managed effectively, protected areas may face consequences that threaten their ability to support both species conservation and the tourism and recreation industries (Rastogi et al., 2015).

Because guest volume is a key driver of HWIs (Cui et al., 2021; Geng, 2021), it is important to understand the factors that shape attendance, as well as how these might alter future attendance. Attendance varies predictably with season and is sensitive to weather, as prevailing weather conditions have a strong impact on the appeal of visiting a protected area and the types of outdoor activities that park-goers participate in (Albano et al., 2013; Fisichelli et al., 2015; Hadwen et al., 2011; Hewer et al., 2016). Protected areas thus have busy seasons, transitory ‘shoulder’ seasons, and off seasons (Butler, 2001), which are primarily determined by temperature and precipitation (Hewer et al., 2016), in combination with the activities available within the protected area. For instance, areas known for skiing will be busiest in the winter, while those used mainly for day hiking or swimming will be busiest in the summer. However, as weather conditions are altered by climate change, such seasonal activities, could be jeopardised and attendance rates may vary considerably (Nyaupane & Chhetri, 2009). The magnitude of these changes will likely be compounded by population growth, since protected areas are likely to experience a greater demand for outdoor recreation and tourism as the human population rise (Balmford et al., 2009). It is thus essential to understand how climate change and population growth will interact to shape visitor counts, HWI rates, and the sustainability of recreation and tourism in protected areas.

The aim of this study is to provide an assessment of the impacts of future climate change and population growth on park tourism in British Columbia, Canada. The Canadian province of British Columbia (BC) is well known for its iconic outdoor recreation industry, which attracts local and international tourists alike and contributes more than US$11 billion in economic value to British Columbians annually (Lloyd-Smith, 2021). Consequently, population growth and climate change are likely to have a profound impact on the province’s income and the sustainability of outdoor recreation and nature-based tourism. All provincially protected areas in British Columbia are managed by BC Parks, a governmental organisation that oversees more than 1,000 protected areas, including conservancies, ecological reserves, and provincial parks (BC Government News, 2021) under the dual mandate of conserving ecological diversity while simultaneously promoting nature-based recreation. The recent growth in the popularity of outdoor recreation and nature-based tourism has caused a rapid rise in provincial park visitors (BC Parks, 2018), which, in turn, has been challenging park operations throughout Canada. In Banff National Park, for instance, interactions with black bears (*Ursus americanus*) attracted to the food brought in by park visitors results in high levels of management-induced bear mortality and relocations (Hebblewhite et al., 2003). Similarly, high densities of visitors in BC’s Cathedral Provincial Park have been spurring an increase in mountain goat (*Oreamnos americanus*) interactions, with reports of goats showing threat displays and forcing hikers off of trails (Balyx, 2022; Vayro et al., 2023).

While increasing guest volume and HWIs challenge the sustainability of tourism and recreation in BC’s parks, there has been no investigation into the potential drivers of these trends, nor whether they might be expected to continue. Like other northern regions, BC’s climate is expected to get warmer and wetter over the coming century (T. Wang et al., 2016), which will likely have pronounced impacts on its tourism industry (Gilani & Innes, 2020). Indeed, previous work has suggested that nature-based tourism in Northern regions is likely to benefit from a warming climate (Steiger et al., 2023). Alongside this trend, BC’s population has nearly doubled since 1980 (Government of British Columbia, 2022) and is presumed to continue growing rapidly. BC’s protected areas are therefore likely to experience notable changes in attendance rates and HWI occurrence through the century as the climate changes and the population grows. Our research will serve as a foundation for understanding and minimizing issues of over crowding and human-wildlife conflict, thereby contributing to sustainable tourism within Canada’s most biodiverse province.

## Methods

### Empirical Data

To understand drivers of park attendance, we analyzed nearly 13 years (January 2010 to August 2022) of day-use attendance records from 249 parks spanning all five of BC Parks’ management regions: Northern, South Coast, West Coast, Thompson-Cariboo, and Kootenay-Okanagan (Fig. 1A). While records of camping were also available for these parks, campsites within the province are limited and generally reserved months in advance and are thus unlikely to reflect decisions based on short-term weather conditions. We thus focused solely on trends in day-use attendance. Attendance records were obtained directly from BC Parks. These data were collected by BC Parks operators using a standardised protocol that estimates visitor counts based on vehicle counts, with buses estimated as 40 visitors and regular vehicles counted as 3.5 visitors. These counts were then totalled for each month and then standardised to ‘visitors per 1,000 people’ based on BC population data obtained from Statistics Canada (Statistics Canada, 2019). Instances of missing visitor counts for a month could not be assumed to indicate no park visitors, and so were treated as NA values. The resulting dataset consisted of 16,469 monthly attendance records, detailing an estimated 41,341,600 individual visits to parks in BC.

**Figure 1.**
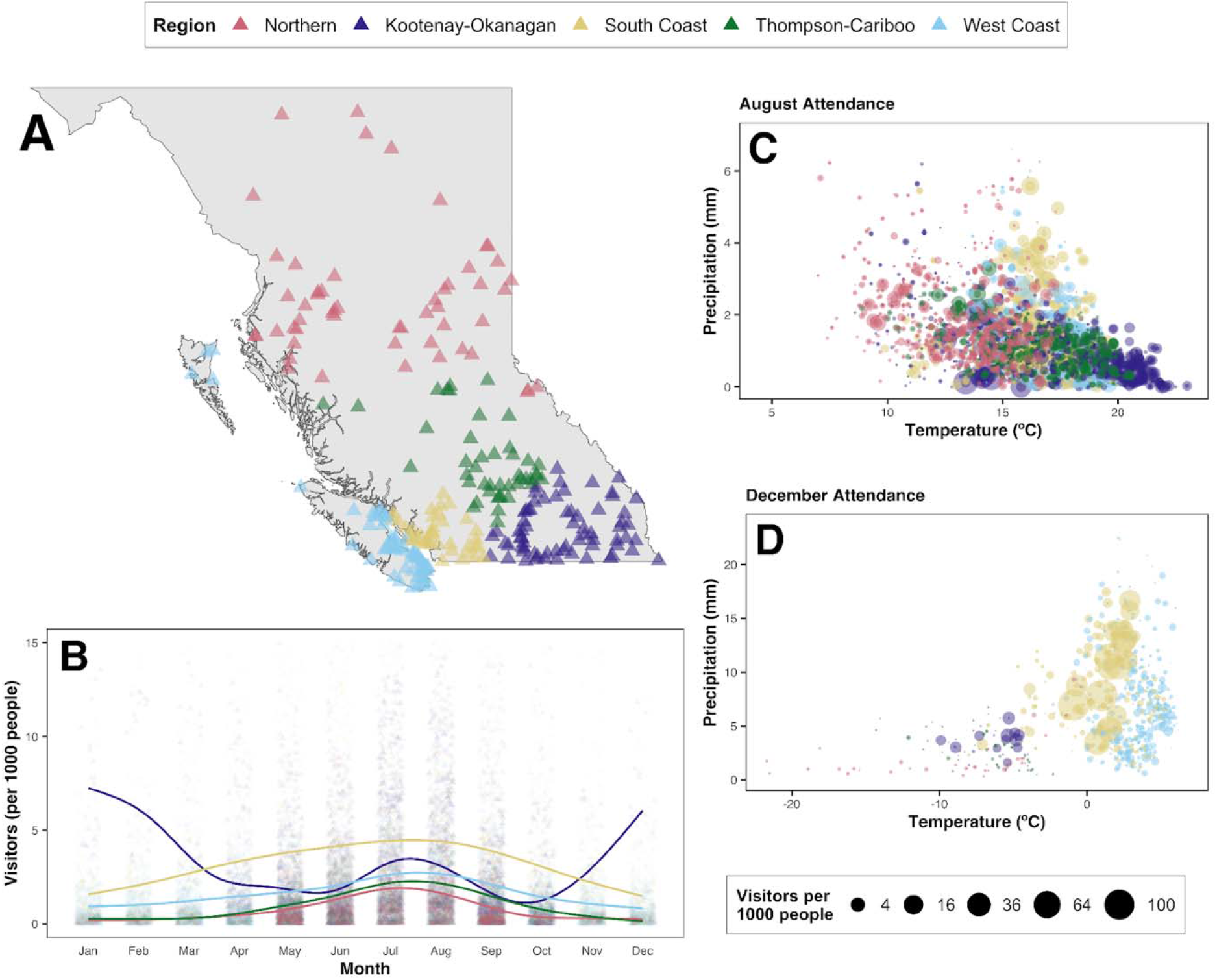
Visualisation of park attendance records. (A) shows the geographic distribution of parks included in this analysis (*n* = 249), coloured by management region. (B) illustrates average seasonal trends for each region from 2010 to 2019. Points represent total monthly visitors, while the smooth lines are average trends across each respective region. (C) demonstrates the relationship between attendance (represented by the size of the points) and weather for August. (D) shows the same relationship but for the month of December. Note that these figures are purely for visualization of the trend lines and do not reflect the interactions later accounted for by our model.

To characterise trends in HWIs, human-wildlife coexistence incidents data were obtained from the Government of Canada’s Open Government database. The Open Data Record dataset is compiled by the Parks Canada Agency and contains records of human-wildlife coexistence incidents from January 2010 to December 2021 for thirty-five national parks and historic sites. Of the 9 incident types in the dataset, we focused solely on records that were classified as ‘Human Wildlife Interactions’ within the five national parks in British Columbia (i.e., Glacier National Park of Canada, Kootenay National Park of Canada, Mount Revelstoke National Park of Canada, Pacific Rim National Park Reserve of Canada, and Yoho National Park of Canada). The individual HWI records were converted to total monthly counts prior to analysis. The resulting dataset consisted of 460 monthly total HWI records describing 4,782 instances of HWI over 136 months (12 years).

Data were cleaned, modelled, and visualized in R (version 4.2.2, R Core Team, 2022). Historical and projected climate data (temperature, precipitation) were obtained from the open source climatenaR R package that accesses data from the ClimateNA software (T. Wang et al., 2016). As the climatenaR package requires the location and elevation data of the study sites, the coordinates for each park were obtained from Google maps, while the associated Digital Elevation Model (DEM) was acquired using the elevatr package (Hollister et al., 2021).

### Modelling Park Attendance and HWIs

To estimate the effects of weather on park attendance, we fit a Hierarchical Generalized Additive Model (HGAM; Pedersen et al., 2019; Wood, 2011) with a gamma distribution and a log link function data using the mgcv R package (version 1.8-41, Wood, 2017). The gamma distribution and log link were selected because the response variable, attendance (visitors per 1,000 people), was continuous and strictly positive. The predictor variables included in this model were smooth terms of month, average monthly temperature (°C), and average monthly precipitation (mm). We also included tensor interaction terms between month and temperature, month and precipitation, and temperature and precipitation, as the effect of temperature and precipitation depends on the month (e.g., people may avoid hiking on a cold and rainy summer day, but they will enjoy skiing with crisp and snowy weather). This model structure allowed us to tease apart the effects of weather on attendance from the seasonal cycle. A cyclic cubic spline was used to account for the cyclical nature of the response variable over the seasons (Fig. 1B). In addition, park ID was included as a random effect to account for differences between parks.

To estimate the effects of weather on HWIs, we fit an HGAM with a Poisson distribution and a log link function using the mgcv package. The Poisson distribution and log link function were selected because the response variable was strictly positive count data. The HWI model included smooth predictors of month, average monthly temperature (°C), and average monthly precipitation (mm). Similarly with our attendance model, we also included tensor interaction terms between month and temperature, month and precipitation, and temperature and precipitation, as the effect of temperature and precipitation depends on the month. Here again, this model structure allowed us to tease apart the effects of weather on HWIs from the seasonal cycle. As with our attendance model, we also included interactions between month and temperature, month and precipitation, and temperature and precipitation. Again, a cyclic cubic spline was used to account for the cyclical nature of the month term, and park was included as a random effect to account for differences between parks.

### Projections Under Climate Change and Population Growth Scenarios

Using the HGAMs, we projected day-use attendance and HWI rates for the current century based on the combined effects of future climate change and population growth. As future climate depends heavily on many unpredictable factors, including global economics and politics, technological improvements, pollution, and energy and land use (van Vuuren et al., 2011), we produced predictions for each of the Intergovernmental Panel on Climate Change (IPCC)’s Shared Socioeconomic Pathways (SSPs; see Chen et al., 2021): SSP 1-2.6 (sustainability), SSP 2-4.5 (‘middle of the road’), SSP 3-7.0 (global inequality), and SSP 5-8.5 (continued fossil fueled based development). Under each of these scenarios, climate change is expected to result in higher temperatures and more precipitation in BC, though the magnitude of these changes is scenario-specific (Wang et al., 2016). As noted above, projected climate data were obtained from the open source ClimateNA software (Wang et al., 2016) via the climatenaR R package.

It is largely assumed that BC’s population will continue to rise. Statistics Canada (Statistics Canada, 2019) predicts BC population growth rates under 9 scenarios (high growth, low growth, medium growth 1 through 5, slow aging, and fast aging) for three years: 2022-2023, 2032-2033, and 2042-2043. We extrapolated growth rates between these decades and beyond to 2100 using a generalized linear mixed-effect model via the R package lme4 (version 1.1-31, Bates et al., 2015). Again, this model had a gamma distribution and a log link function.

According to our population growth model, BC’s population is expected to increase under all population growth scenarios (see Supplementary Material). We focused primarily on the high and low growth scenarios, but projections for all 9 population growth scenarios are openly available on our GitHub repository (link below).

All the data, R scripts, supporting figures, and supplementary material are openly available at https://github.com/QuantitativeEcologyLab/BCParks_Attendance.

## Results

### Weather, Seasons, and Park Attendance

Overall, we found that prevailing weather conditions and season had substantial impacts on park attendance. Attendance was positively correlated with temperature (Fig. 2A), with summer (June through August) being the peak season for most regions (Fig. 1B). There was also a negative relationship between park attendance and precipitation, with heavier rains tending to result in lower attendance, although the effect of precipitation was much weaker than the effect of temperature trends varied (Fig. 2B). Importantly though, the relationship between attendance and weather changed based on the seasonal context, and park attendance was impacted by both the marginal and interaction effects of temperature, precipitation, and time of year. For instance, parks tended to receive fewer visitors in unusually warm summers, but they received more visitors during unusually warm winters (see e.g., Fig. 1C, and 1D).

**Figure 2.**
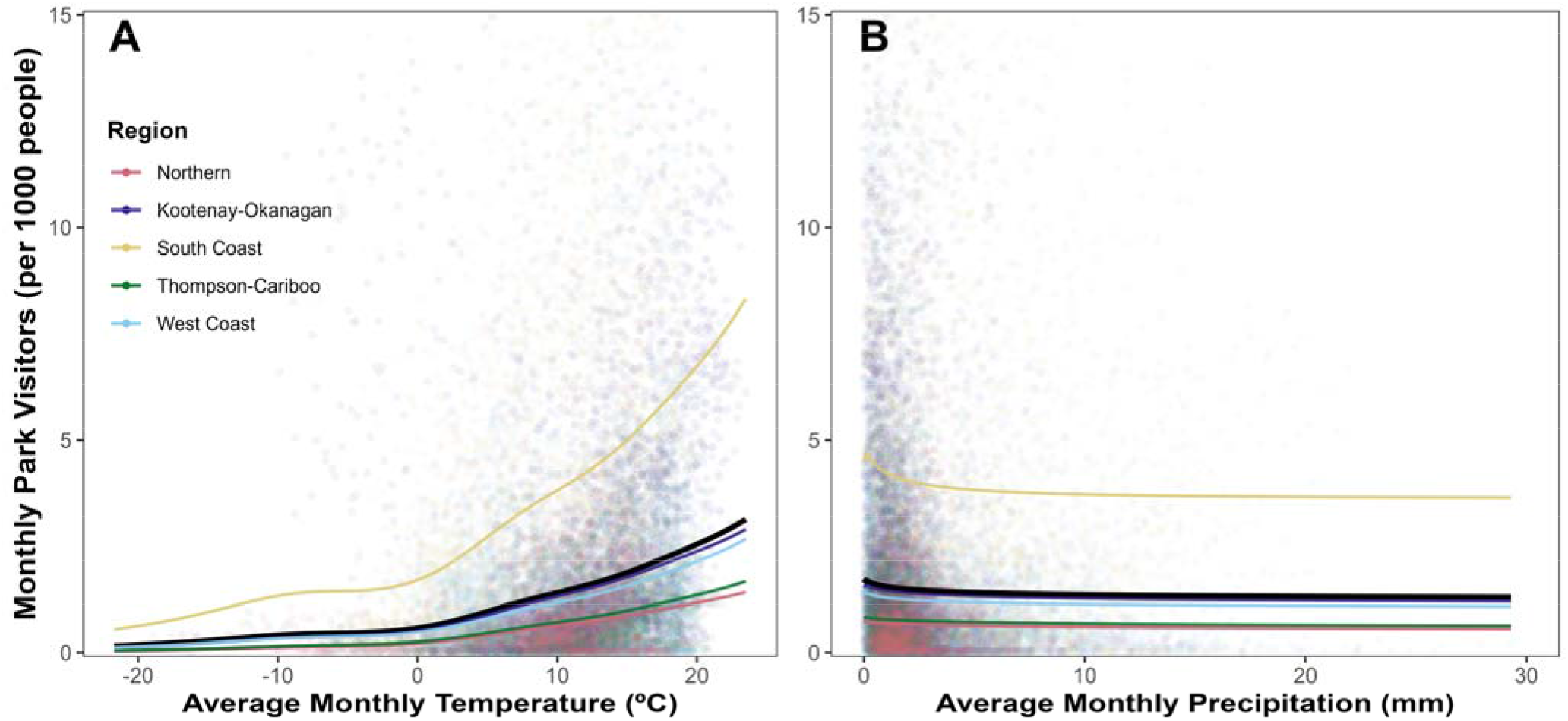
Relationship between BC Parks attendance and weather. Observations are represented by points, while the coloured lines show the model-predicted trends in regional attendance. (A) shows a positive, non-linear relationship between the number of visitors and average monthly temperature overall, although regions differ in exact trends. (B) shows that increasing precipitation generally has a negative, though relatively weak impact on attendance.

### Climate Change and Projected Attendance

Due to the strong relationships between park attendance and weather conditions, visitation was generally projected to increase under each of the climate change scenarios (Fig. 3). However, the magnitude of increase differed strongly between scenarios. Under SSP 1-2.6 (the more optimal pathway), attendance rates were little changed, with only a ca. 20% increase in attendance. In contrast, attendance was projected to nearly double under the most drastic scenario (SSP 5-8.5).

**Figure 3.**
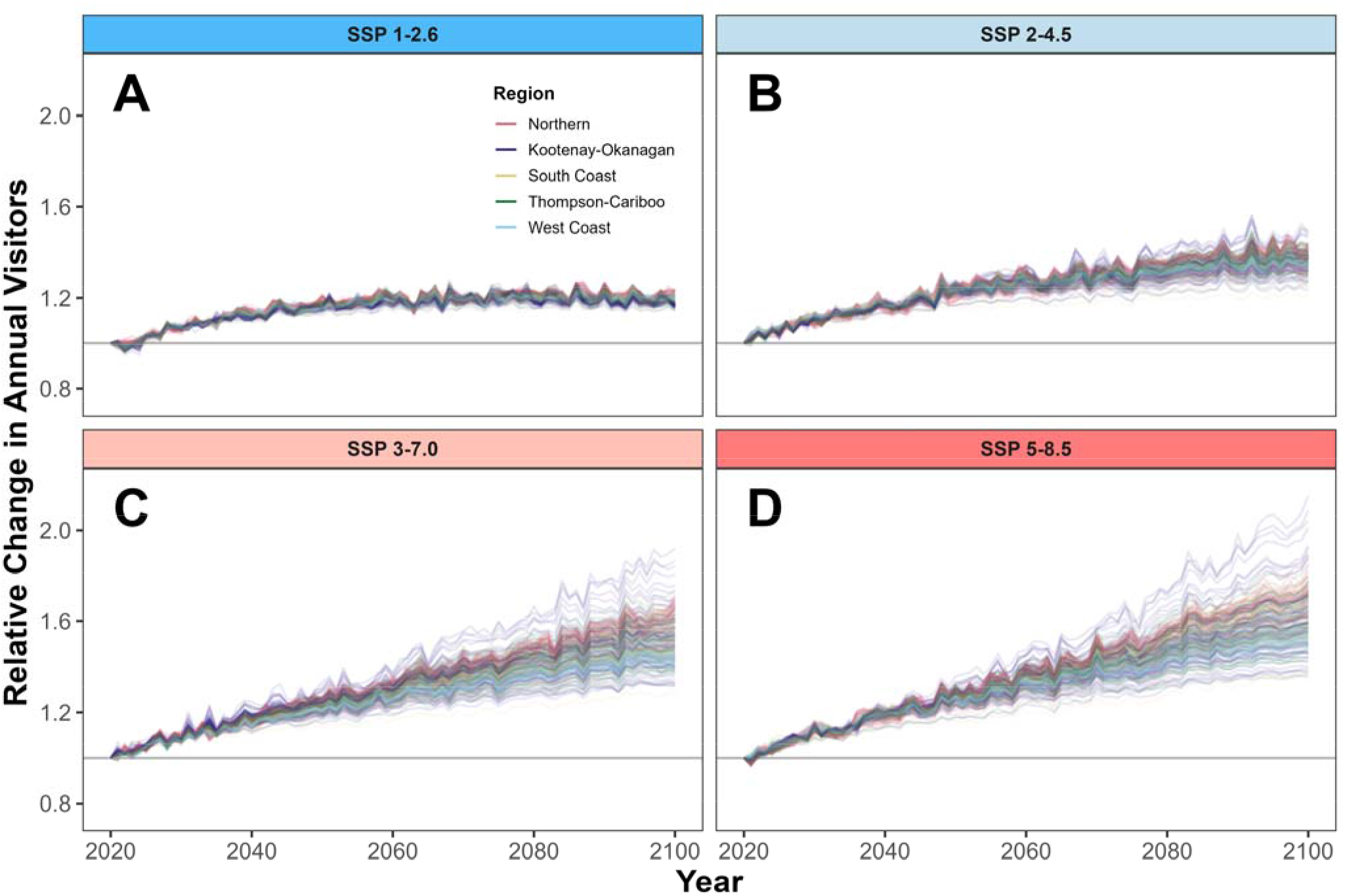
Attendance projections for 249 parks. Each line represents one park’s change in visitors relative to 2020. Projection lines are coloured by region and are presented for the four climate change scenarios corresponding to each Shared Socioeconomic Pathway (SSP) hypothesized by the IPCC. The horizontal grey line at 1 indicates no relative change in attendance. These panels exclusively show the effect of climate change on park attendance; the effect of population growth is not visualized here.

#### Case Study: Golden Ears Park

The results presented above serve as an overview of day-use park attendance in the upcoming century. Notably though, the values shown in figure 3 are on a relative scale, and the absolute number of visitors will also depend on population growth. To examine how the model projected seasonal trends under different combinations of climate change and population growth, we investigated Golden Ears Park as a case study. Our projections suggest that, under any combination of scenarios, Golden Ears Park will experience a substantial rise in visitors moving forward (Fig. 4). Consistent with the general trends described above, the lower-emissions climate scenarios (SSP 1-2.6 and 2-4.5) were projected to lead to fewer visitors than the higher-emissions scenarios (SSP 3-7.0 and 5-8.5). Not only did our models project substantially busier peak seasons, but it appears that shoulder seasons will quickly approach and succeed attendance rates comparable to that of the current peak season. High levels of population growth further amplified the effects of climate change on park attendance (Fig. 4B). While we only present results for Golden Ears Park here, similar patterns were observed across all 249 parks (Supplementary Material).

**Figure 4.**
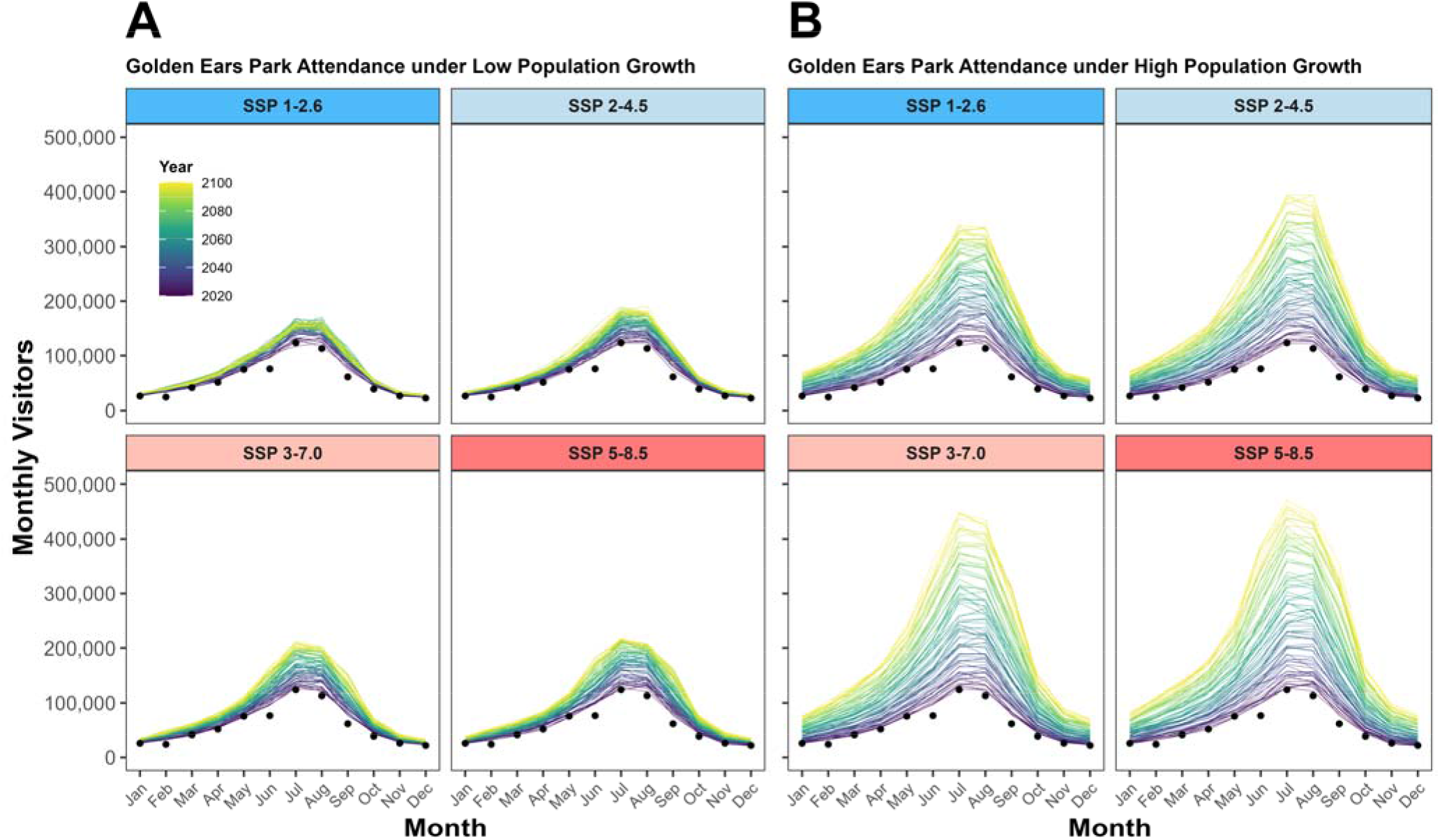
Attendance projections for Golden Ears Park under different population growth and climate change scenarios. Average historical attendance from 2010 to 2019 is represented by black points. Projections are coloured according to year, ranging from 2020 to 2100. Attendance rates depend on the climate change scenario (facets; SSP 1-2.6, SSP 2-4.5, SSP 3-7.0, and SSP 5-8.5) and the population growth scenario. (A) shows projected attendance when BC is under a low population growth scenario, and (B) shows projected attendance for a high population growth scenario.

### The Future of Human-Wildlife-Interactions

When modelling HWIs, we found that the same weather conditions that drove more people to visit parks also resulted in greater HWI rates. The number of HWIs was positively correlated with temperature (Fig. 5A), and there was also a negative (albeit weaker) relationship between HWIs and precipitation (Fig. 5B). Consequently, HWIs were projected to increase under all four climate change scenarios we investigated (Fig. 5C-F). As with park attendance, we found markedly different outcomes depending on which climate change scenario we assumed would occur. Under the lower emissions scenarios like SSPs 1-2.6, our model projected a minor (ca. 10%) increase in HWIs. In contrast, the ‘worst-case’ climate change scenario (SSP 5-8.5) is expected to result in a substantial increase in the number of HWIs, though projected trends still differ between parks.

**Figure 5.**
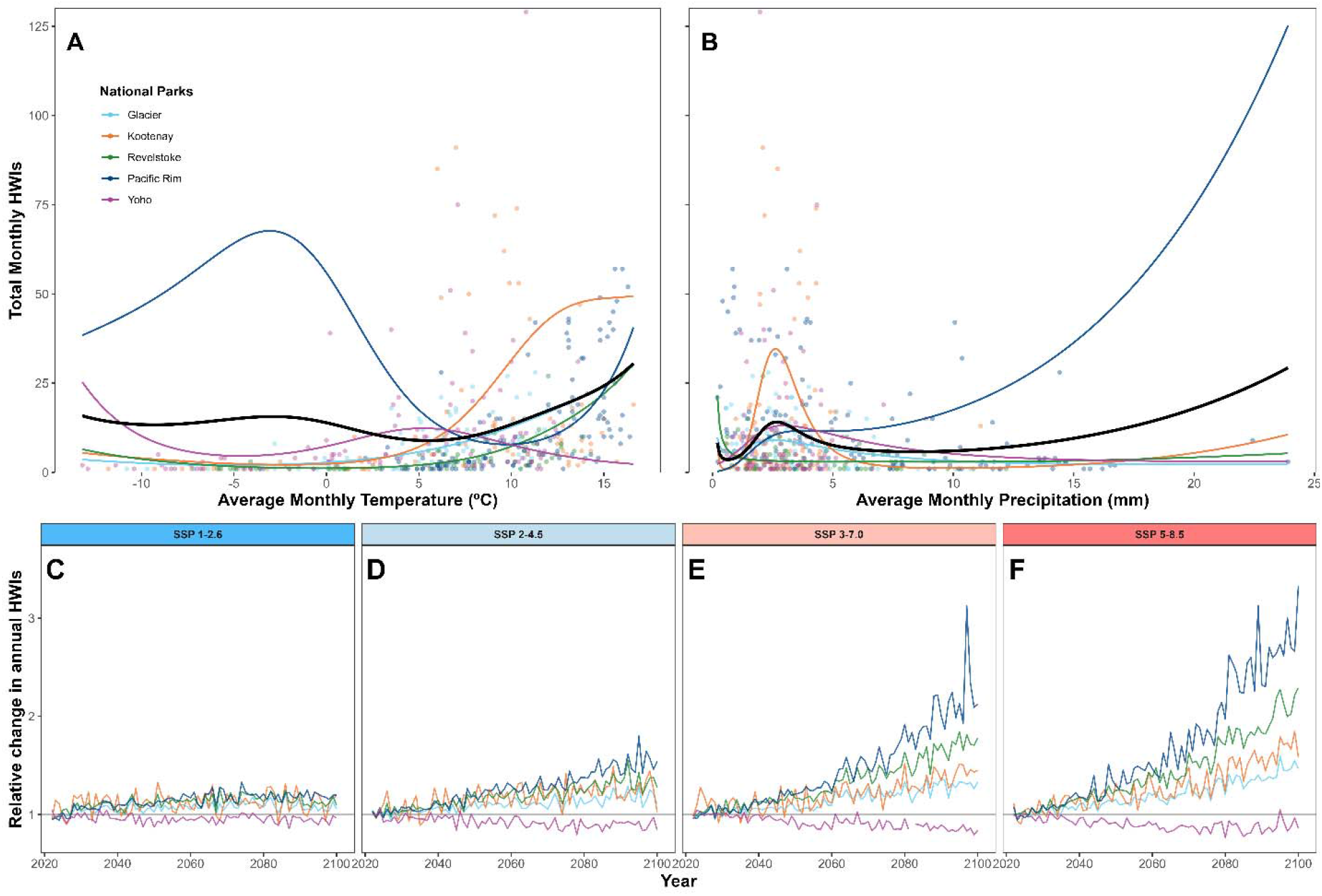
Effects of temperature and precipitation on human-wildlife interactions and projections for the current century in BC national parks. The top row shows the relationship between the number of human-wildlife interactions (HWIs) and average monthly (A) temperature and (B) precipitation. Observed HWIs are represented by points, while thin coloured lines show park specific trends and overall trends are represented by thick, black lines. The bottom row depicts the projected relative change in HWIs for these parks. Each line represents one park’s annual change in HWIs relative to 2022 for the four climate change scenarios. The horizontal grey line at 1 indicates no relative change in annual HWIs.

## Discussion

This study sought to model and project trends in attendance and human-wildlife-interactions in British Columbia’s protected areas through to the end of the next century. This work allows park managers to make proactive management decisions that will help ensure the sustainability of nature-based tourism within the province. Our findings are consistent with recent work by (Steiger et al., 2023), who found that nature-based tourism in Northern regions, such as British Columbia, is likely to benefit from a warming climate. Notably though, increased attendance is not necessarily a beneficial outcome, as high volumes of guests can strain operations (Manning, 2001) and negatively impact wildlife (Cui et al., 2021; Geng, 2021). Indeed, we found that the same weather conditions that drove more people to visit parks also resulted in more HWIs, highlighting the intrinsic relationship between weather, park attendance, and HWIs. If not properly managed for, climate change-driven increases in attendance and HWIs risk challenging the sustainability of nature-based tourism within British Columbia’s network of protected areas.

The tight correlation between attendance and weather was not surprising. It is well recognised that weather shapes the appeal of visiting a protected area and the types of outdoor activities that park-goers will participate in (Albano et al., 2013; Fisichelli et al., 2015; Hadwen et al., 2011; Hewer et al., 2016). Importantly, however, we were able to tease apart the effects of weather from the seasonal cycle (Hadwen et al., 2011) and leverage these relationships to project attendance and HWIs through to the end of the century. While there is debate as to which climate change scenario is considered the most likely outcome based on past and current trends (Hausfather & Peters, 2020; Schwalm et al., 2020), the goal of our work was not to assess the probability of each pathway, so we provide predictions for each of the SSPs. It is also important to note that we projected total expected HWIs across all species, yet there will likely be interspecific differences in HWIs. Black bears and mountain goats, for instance, are often attracted towards people and HWIs may increase for these species if animals are rewarded with resources or predator shields (Balyx, 2022; Hebblewhite et al., 2003). Other species such as wolves (*Canis lupus*) or caribou (*Rangifer tarandus tarandus*) tend to avoid or change their behaviour in areas where humans recreate (Gill et al., under review; Rogala et al., 2011), which would result in a reduction in HWIs with rising park attendance. Future research should aim to identify the effects of increasing visitor volumes across species (e.g., Procko et al., 2023).

Rising attendance rates and HWIs have the potential to severely disrupt the day-to-day operations of protected areas, particularly in the case of highly crowded parks, and threaten the sustainability of nature-based tourism (Stem et al., 2003; Stronza et al., 2019). While increasing the number of parking spaces, trails, campsites, and guest facilities may mitigate operational challenges caused by overcrowding, these solutions can result in a loss of conservation value in parks (Jones et al., 2018; Sarmento & Berger, 2017; Stronza et al., 2019), violating the dual mandate that BC Parks and similar organizations operate under. To mitigate these challenges, we recommend that park managers make proactive decisions based on projections such as those presented here, in order to be better prepared for the increase in attendance and HWIs they are likely to experience over the coming decades. In the near term, an increase in staff may help alleviate the pressure of additional visitors (including substantial increases during shoulder seasons). Longer term, however, managers should carefully evaluate their seasonal carrying capacities (Manning, 2001) and consider imposing guest limits. Yet rather than taking a one-size-fits-all approach, managers should make decisions on a park-by-park basis (Baker, 1992), and actively consider how higher attendance rates will affect each unique park and what further action must be taken to comply with BC Parks’ dual mandate. To assist in understanding individual park trends, we generated detailed projections for all 249 parks (available in our public GitHub repository; see the data availability statement).

Though our models were able to accurately capture trends in the historical data, there are many consequences of climate change that could influence attendance but are not accounted for in our predictions. For instance, our model did not account for thermal thresholds at which people might lose interest in visiting a park, whether it be because temperatures are too high or too low (Fisichelli et al., 2015). However, research by (Hewer et al., 2016) suggests that parks may still experience a rise in annual visitors despite exceeding a maximum thermal threshold. Extreme climatic events such as heat domes, wildfires, and flash floods also have the potential to severely disrupt tourism and recreation in protected areas (Nyaupane & Chhetri, 2009). Although it was beyond the scope of this study to make predictions about the future occurrence of natural disasters, the frequency of these events is expected to increase in the current century (X. Wang et al., 2015), so managers should strategize methods of managing protected areas based on parks’ unique attributes (Baker, 1992). Additionally, it is worth noting that our projections were based on average monthly predictions of future temperature and precipitation. Consequently, our predictions do not account for weather variations within months, including short-term extreme weather events. Changes in snow cover (and changes in the frequency of avalanches) are also expected to affect park attendance, but we did not include snow data in the present study due to a lack of snow cover projection models.

In addition to extreme weather events, climate change is also predicted to have ecological consequences that could affect park attendance. Wildlife is a major attraction for some BC Parks, and peak seasons at these parks align with the timing of annual ecological events. For instance, Rathtrevor Beach Park is often attended for bird-watching opportunities when seabirds gather to feed on spawning herrings in spring and Brant geese migrate in late winter (BC Parks, 2023b). In contrast, the busiest season at Kokanee Creek Park is mid-August to mid-September, as this is when visitors visit the park to see salmon spawning in the channels (BC Parks, 2023a). As the climate changes, wildlife may shift the timing of reproduction or migration (Cadahía et al., 2017; Matthysen et al., 2011; Moyes et al., 2010), which will likely change the timing of peak months. Further research might thus explore local phenology in response to climate change to predict the future peak seasons in parks with wildlife attractions.

## Conclusion

This study provides insight into the patterns of park attendance and HWIs during the coming century, along with a valuable framework for modelling and predicting the two. We have demonstrated that both park attendance and HWI frequency will increase in the next few decades, regardless of which climate change or population growth scenarios occur. Consequently, proactive management is urgently required by BC Parks. Without adequate planning, human-wildlife conflict will likely escalate, which will challenge BC Parks’ dual mandate and render recreation in these parks unsustainable. Park managers should use these findings to plan for more visitors and avoid cases of human-wildlife conflict, ultimately sustaining the viability of nature-based tourism.

## Funding details

This work was supported by an NSERC Discovery Grant RGPIN-2021-02758, the Canadian Foundation for Innovation, as well as the BC Parks Living Labs funding program.

## Disclosure statement

Authors declare that they have no competing interests.

## Biographical note

Dayna Weststrate completed her BSc Hons in Biology at the University of British Columbia Okanagan. She is pursuing medicine in her home province of British Columbia, though her passion for conservation and sustainability remains. Aimee Chhen is a researcher in the Quantitative Ecology Lab. She has a BSc in Zoology and an interest in animal behaviour and species conservation. Stefano Mezzini is a PhD student at the University of British Columbia Okanagan studying how environmental unpredictability affects how often, how much, and where animals move. He completed his BSc Hons in Biology and BSc in Statistics at the University of Regina (SK), and he is interested in understanding the large-scale processes that drive animal movement and behavior. Kirk Safford is a conservation specialist for the Ministry of Environment and Climate Change Strategy. He works primarily within the Kootenay Okanagan Region on applied challenges, including mitigating human-wildlife interactions.

Michael Noonan is a quantitative ecologist and head of the Quantitative Ecology Lab at the University of British Columbia Okanagan. Noonan’s research is focused on protecting vulnerable species by using data-driven approaches to ensure that the evidence used to support evidence based conservation is both accurate and reliable.

## Data availability

The data and R scripts used to carry out this study are openly available on GitHub at https://github.com/QuantitativeEcologyLab/BCParks_Attendance.

## Notes

https://github.com/QuantitativeEcologyLab/BCParks_Attendance

### Competing Interest Statement

The authors have declared no competing interest.

## References

Albano, C. M., Angelo, C. L., & Thurman, L. L. (2013). Potential effects of warming climate on visitor use in three Alaskan national parks. Park Science, 30(1), 36–44.

Baker, W. L. (1992). The landscape ecology of large disturbances in the design and management of nature reserves. Landscape Ecology, 7(3), 181–194. https://doi.org/10.1007/BF00133309

Balmford, A., Beresford, J., Green, J., Naidoo, R., Walpole, M., & Manica, A. (2009). A Global Perspective on Trends in Nature-Based Tourism. PLoS Biology, 7, e1000144. https://doi.org/10.1371/journal.pbio.1000144

Balyx, L. (2022). Human conflict and coexistence with mountain goats in a protected alpine landscape (Version 1) [MSc, University of British Columbia]. https://dx.doi.org/10.14288/1.0416294

Bass, M. S., Finer, M., Jenkins, C. N., Kreft, H., Cisneros-Heredia, D. F., McCracken, S. F., Pitman, N. C. A., English, P. H., Swing, K., Villa, G., Di Fiore, A., Voigt, C. C., & Kunz, T. H. (2010). Global conservation significance of Ecuador’s Yasuní National Park. PloS One, 5(1), e8767. https://doi.org/10.1371/journal.pone.0008767

Bates, D., Maechler, M., Bolker, B., & Walker, S. (2015). Fitting Linear Mixed-Effects Models Using lme4. Journal of Statistical Software, 67(1), 1–48. https://doi.org/10.18637/jss.v067.i01

BC Government News. (2021, January 13). Province acquires land for 16 provincial parks, two protected areas. Environment and Climate Change Strategy. https://news.gov.bc.ca/releases/2021ENV0001-000029

BC Parks. (2018). BC Parks 2017/18 Statistics Report (p. 6). https://nrs.objectstore.gov.bc.ca/kuwyyf/statistic_report_2017_2018_20845e2e6c.pdf

BC Parks. (2023a). Kokanee Creek Park [Government]. BC Parks. https://bcparks.ca/kokanee-creek-park/

BC Parks. (2023b). Rathtrevor Beach Park [Government]. BC Parks. https://bcparks.ca/rathtrevor-beach-park/

Buckley, R. (2018). Tourism and Natural World Heritage: A Complicated Relationship. Journal of Travel Research, 57(5), 563–578. https://doi.org/10.1177/0047287517713723

Buckley, R., Brough, P., Hague, L., Chauvenet, A., Fleming, C., Roche, E., Sofija, E., & Harris, N. (2019). Economic value of protected areas via visitor mental health. Nature Communications, 10(1), 5005. https://doi.org/10.1038/s41467-019-12631-6

Butler, R. W. (2001). Seasonality in Tourism: Issues and Implications. In T. Baum & S. Lundtorp (Eds.), Seasonality in Tourism (1st Edition, pp. 5–21). Routledge.

Cadahía, L., Labra, A., Knudsen, E., Nilsson, A., Lampe, H. M., Slagsvold, T., & Stenseth, N. C. (2017). Advancement of spring arrival in a long-term study of a passerine bird: Sex, age and environmentalo effects. Oecologia, 184, 917–929. https://doi.org/0.1007/s00442-017-3922-4

Charron, D. (2002). Potential impacts of global warming and climate change on the epidemiology of zoonotic diseases in Canada. Canadian Journal of Public Health, 93(5), 334.

Chen, C., Brodie, J. F., Kays, R., Davies, T. J., Liu, R., Fisher, J. T., Ahumada, J., McShea, W., Sheil, D., & Agwanda, B. (2022). Global camera trap synthesis highlights the importance of protected areas in maintaining mammal diversity. Conservation Letters, 15(2), e12865.

Chen, D., Rojas, M., Samset, B. H., Cobb, K., Diongue Niang, A., Edwards, P., Emori, S., Faria, S. H., Hawkins, E., Hope, P., Huybrechts, P., Meinshausen, M., Mustafa, S. K., Plattner, G.-K., & Tréguier, A.-M. (2021). 2021: Framing, Context, and Methods. In M.-D. V., P. Zhai, A. Pirani, S. L. Connors, C. Péan, S. Berger, N. Caud, Y. Chen, L. Goldfarb, M. I. Gomis, M. Huang, K. Leitzell, E. Lonnoy, J. B. R. Matthews, T. K. Maycock, T. Waterfield, O. Yelekçi, R. Yu, & B. Zhou (Eds.), Climate Change 2021: The Physical Science Basis. Contribution of Working Group I to the Sixth Assessment Report of the Intergovernmental Panel on Climate Change (pp. 147–286). Cambridge University Press. https://www.ipcc.ch/report/ar6/wg1/chapter/chapter-1/

Corradini, A., Randles, M., Pedrotti, L., van Loon, E., Passoni, G., Oberosler, V., Rovero, F., Tattoni, C., Ciolli, M., & Cagnacci, F. (2021). Effects of cumulated outdoor activity on wildlife habitat use. Biological Conservation, 253, 108818. https://doi.org/10.1016/j.biocon.2020.108818

Cui, Q., Ren, Y., & Xu, H. (2021). The Escalating Effects of Wildlife Tourism on Human Wildlife Conflict. AnimalslJ: An Open Access Journal from MDPI, 11(5). https://doi.org/10.3390/ani11051378

Eagles, P. F. J. (2002). Trends in Park Tourism: Economics, Finance and Management. Journal of Sustainable Tourism, 10(2), 132–153. https://doi.org/10.1080/09669580208667158

Fisichelli, N. A., Schuurman, G. W., Monahan, W. B., & Ziesler, P. S. (2015). Protected Area Tourism in a Changing Climate: Will Visitation at US National Parks Warm Up or Overheat? PLOS ONE, 10(6), e0128226. https://doi.org/10.1371/journal.pone.0128226

Gaynor, K. M., Hojnowski, C. E., Carter, N. H., & Brashares, J. S. (2018). The influence of human disturbance on wildlife nocturnality. Science, 360(6394), 1232–1235. https://doi.org/10.1126/science.aar7121

Geng, D. (2021). Managing national park visitor experience and visitor-wildlife coexistence: A case study of Banff National Park [Text, University of British Columbia]. https://open.library.ubc.ca/collections/24/items/1.0398303

Gilani, H. R., & Innes, J. L. (2020). The State of British Columbia’s Forests: A Global Comparison. Forests, 11(3). https://doi.org/10.3390/f11030316

Gill, R., Serouya, R., Ford, A., Steenweg, R., Calvert, A. M., & Noonan, M. J. (under review). Movement Ecology of Endangered Caribou During a COVID-19 Mediated Pause in Winter Recreation. Animal Conservation.

Government of British Columbia. (2022). Annual population, July 1, 1867-2022.

Gray, C. L., Hill, S. L. L., Newbold, T., Hudson, L. N., Börger, L., Contu, S., Hoskins, A. J., Ferrier, S., Purvis, A., & Scharlemann, J. P. W. (2016). Local biodiversity is higher inside than outside terrestrial protected areas worldwide. Nature Communications, 7(1), 12306. https://doi.org/10.1038/ncomms12306

Hadwen, W. L., Arthington, A. H., Boon, P. I., Taylor, B., & Fellows, C. S. (2011). Do Climatic or Institutional Factors Drive Seasonal Patterns of Tourism Visitation to Protected Areas across Diverse Climate Zones in Eastern Australia? Tourism Geographies, 13(2), 187–208. https://doi.org/10.1080/14616688.2011.569568

Hausfather, Z., & Peters, G. P. (2020). Emissions—The “business as usual” story is misleading. Nature, 577(7792), 618–620. https://doi.org/10.1038/d41586-020-00177-3

Hebblewhite, M., Percy, M., & Serrouya, R. (2003). Black bear (Ursus americanus) survival and demography in the Bow Valley of Banff National Park, Alberta. Biological Conservation, 112(3), 415–425. https://doi.org/10.1016/S0006-3207(02)00341-5

Hewer, M., Scott, D., & Fenech, A. (2016). Seasonal weather sensitivity, temperature thresholds, and climate change impacts for park visitation. Tourism Geographies, 18(3), 297–321. https://doi.org/10.1080/14616688.2016.1172662

Hollister, J., Shah, T., Robitaille, A. L., Beck, M. W., & Johnson, M. (2021). elevatr: Access elevation data from various APIs [Manual]. https://doi.org/10.5281/zenodo.5809645

Jones, K. R., Venter, O., Fuller, R. A., Allan, J. R., Maxwell, S. L., Negret, P. J., & Watson, J. E. M. (2018). One-third of global protected land is under intense human pressure. Science, 360(6390), 788–791. https://doi.org/10.1126/science.aap9565

Larson, C. L., Reed, S. E., Merenlender, A. M., & Crooks, K. R. (2016). Effects of Recreation on Animals Revealed as Widespread through a Global Systematic Review. PLOS ONE, 11(12), e0167259. https://doi.org/10.1371/journal.pone.0167259

Lloyd-Smith, P. (2021). The economic benefits of recreation in Canada. *Canadian Journal of Economics/Revue Canadienne d’**é*conomique, 54(4), 1684–1715. https://doi.org/10.1111/caje.12560

Lopez Gutierrez, B., Almeyda Zambrano, A. M., Mulder, G., Ols, C., Dirzo, R., Almeyda Zambrano, S. L., Quispe Gil, C. A., Cruz Díaz, J. C., Alvarez, D., Valdelomar Leon, V., Villareal, E., Sanchez Espinosa, A., Quiros, A., Stein, T. V., Lewis, K., & Broadbent, E. N. (2020). Ecotourism: The ‘human shield’ for wildlife conservation in the Osa Peninsula, Costa Rica. Journal of Ecotourism, 19(3), 197–216. https://doi.org/10.1080/14724049.2019.1686006

Manning, R. (2001). Visitor experience and resource protection: A framework for managing the carrying capacity of National Parks. Journal of Park & Recreation Administration, 19(1), 93–108.

Margules, C. R., & Pressey, R. L. (2000). Systematic conservation planning. Nature, 405(6783), 243–253. https://doi.org/10.1038/35012251

Matthysen, E., Adriaensen, F., & Dhondt, A. A. (2011). Multiple responses to increasing spring temperatures in the breeding cycle of blue and great tits (Cyanistes caeruleus, Parus major). Global Change Biology, 17(1), 1–16. https://doi.org/10.1111/j.1365-2486.2010.02213.x

Monahan, A. M., Miller, I. S., & Nally, J. E. (2009). Leptospirosis: Risks during recreational activities. Journal of Applied Microbiology, 107(3), 707–716. https://doi.org/10.1111/j.1365-2672.2009.04220.x

Moyes, K., Nussey, D. H., Clements, M. N., Guinness, F. E., Morris, A., Morris, S., Pemberton, J. M., Kruuk, L. E., & Clutton-Brock, T. H. (2010). Advancing breeding phenology in response to environmental change in a wild red deer population. Global Change Biology, 17(7), 2455–2469. https://doi.org/10.1111/j.1365-2486.2010.02382.x

Nyaupane, G. P., & Chhetri, N. (2009). Vulnerability to Climate Change of Nature-Based Tourism in the Nepalese Himalayas. Tourism Geographies, 11(1), 95–119. https://doi.org/10.1080/14616680802643359

Nyhus, P. J. (2016). Human–Wildlife Conflict and Coexistence. Annual Review of Environment and Resources, 41(1), 143–171. https://doi.org/10.1146/annurev-environ-110615-085634

Pedersen, E. J., Miller, D. L., Simpson, G. L., & Ross, N. (2019). Hierarchical generalized additive models in ecology: An introduction with mgcv. PeerJ, 7, e6876. https://doi.org/10.7717/peerj.6876

Prakash, S. L., Perera, P., Newsome, D., Kusuminda, T., & Walker, O. (2019). Reasons for visitor dissatisfaction with wildlife tourism experiences at highly visited national parks in Sri Lanka. Journal of Outdoor Recreation and Tourism, 25, 102–112. https://doi.org/10.1016/j.jort.2018.07.004

Procko, M., Naidoo, R., LeMay, V., & Burton, A. C. (2023). Human presence and infrastructure impact wildlife nocturnality differently across an assemblage of mammalian species. PLOS ONE, 18(5), e0286131. https://doi.org/10.1371/journal.pone.0286131

R Core Team. (2022). R: A language and environment for statistical computing (4.2.2) [R]. https://www.R-project.org/

Rastogi, A., Hickey, G. M., Anand, A., Badola, R., & Hussain, S. A. (2015). Wildlife-tourism, local communities and tiger conservation: A village-level study in Corbett Tiger Reserve, India. Forest Policy and Economics, 61, 11–19. https://doi.org/10.1016/j.forpol.2015.04.007

Reed, S. E., & Merenlender, A. M. (2008). Quiet, Nonconsumptive Recreation Reduces Protected Area Effectiveness: Quiet recreation in protected areas. Conservation Letters, 1(3), 146–154. https://doi.org/10.1111/j.1755-263X.2008.00019.x

Rogala, J. K., Hebblewhite, M., Whittington, J., White, C. A., Coleshill, J., & Musiani, M. (2011). Human Activity Differentially Redistributes Large Mammals in the Canadian Rockies National Parks. Ecology and Society, 16(3), art16. https://doi.org/10.5751/ES-04251-160316

Sangpikul, A. (2017). Ecotourism Impacts on the Economy, Society and Environment of Thailand. Journal of Reviews on Global Economics, 6, 302–312. https://doi.org/10.6000/1929-7092.2017.06.30

Sarmento, W. M., & Berger, J. (2017). Human visitation limits the utility of protected areas as ecological baselines. Biological Conservation, 212, 316–326. https://doi.org/10.1016/j.biocon.2017.06.032

Schwalm, C. R., Glendon, S., & Duffy, P. B. (2020). RCP8.5 tracks cumulative CO(2) emissions. Proceedings of the National Academy of Sciences of the United States of America, 117(33), 19656–19657. https://doi.org/10.1073/pnas.2007117117

Statistics Canada. (2019). Population Projections for Canada (2018-2068), Provinces and Territories (2018 to 2043). Statistics Canada. https://www150.statcan.gc.ca/n1/en/pub/91-520-x/91-520-x2019001-eng.pdf?st=VkaTtamd

Steiger, R., Demiroglu, O. C., Pons, M., & Salim, E. (2023). Climate and carbon risk of tourism in Europe. Journal of Sustainable Tourism, 1–31. https://doi.org/10.1080/09669582.2022.2163653

Stem, C. J., Lassoie, J. P., Lee, D. R., & Deshler, D. J. (2003). How “Eco” is Ecotourism? A Comparative Case Study of Ecotourism in Costa Rica. Journal of Sustainable Tourism, 11(4), 322–347. https://doi.org/10.1080/09669580308667210

Stronza, A. L., Hunt, C. A., & Fitzgerald, L. A. (2019). Ecotourism for Conservation? Annual Review of Environment and Resources, 44(1), 229–253. https://doi.org/10.1146/annurev-environ-101718-033046

Taylor, J. E., Dyer, G. A., Stewart, M., Yunez_JNaude, A., & Ardila, S. (2003). The Economics of Ecotourism: A Galápagos Islands Economy_JWide Perspective. Economic Development and Cultural Change, 51(4), 977–997. https://doi.org/10.1086/377065

Thomas, C. D., & Gillingham, P. K. (2015). The performance of protected areas for biodiversity under climate change. Biological Journal of the Linnean Society, 115(3), 718–730. https://doi.org/10.1111/bij.12510

van Vuuren, D. P., Edmonds, J., Kainuma, M., Riahi, K., Thomson, A., Hibbard, K., Hurtt, G. C., Kram, T., Krey, V., Lamarque, J.-F., Masui, T., Meinshausen, M., Nakicenovic, N., Smith, S. J., & Rose, S. K. (2011). The representative concentration pathways: An overview. Climatic Change, 109(1), 5. https://doi.org/10.1007/s10584-011-0148-z

Vayro, J. V., Vandermale, E. A., & Mason, C. W. (2023). ‘It’s a people problem, not a goat problem.’ Mitigating human–mountain goat interactions in a Canadian Provincial Park. Wildlife Research. https://doi.org/10.1071/WR22005

Wang, T., Hamann, A., Spittlehouse, D., & Carroll, C. (2016). Locally Downscaled and Spatially Customizable Climate Data for Historical and Future Periods for North America. PLOS ONE, 11(6), e0156720. https://doi.org/10.1371/journal.pone.0156720

Wang, X., Thompson, D. K., Marshall, G. A., Tymstra, C., Carr, R., & Flannigan, M. D. (2015). Increasing frequency of extreme fire weather in Canada with climate change. Climatic Change, 130(4), 573–586. https://doi.org/10.1007/s10584-015-1375-5

Whittington, J., Hebblewhite, M., Baron, R. W., Ford, A. T., & Paczkowski, J. (2022). Towns and trails drive carnivore movement behaviour, resource selection, and connectivity. Movement Ecology, 10(1), 17. https://doi.org/10.1186/s40462-022-00318-5

Wood, S. N. (2011). Fast Stable Restricted Maximum Likelihood and Marginal Likelihood Estimation of Semiparametric Generalized Linear Models. Journal of the Royal Statistical Society Series B: Statistical Methodology, 73(1), 3–36. https://doi.org/10.1111/j.1467-9868.2010.00749.x

Wood, S. N. (2017). Generalized Additive Models: An Introduction with R (2nd ed.). Chapman and Hall/CRC. https://doi.org/10.1201/9781315370279

